# Application of D4 Fluorescent Probes for Quantitative and Spatial Analysis of Cholesterol in Cells

**DOI:** 10.64898/2026.04.01.715848

**Authors:** Aliénor Passerat de La Chapelle, Elizaveta Boiko, Cem Karakus, Alice Trahin, Anaïs Aulas, Coralie Di Scala

## Abstract

Cholesterol is a key component of cellular membranes, regulating membrane organization, fluidity, and signaling. However, cholesterol analysis remains technically challenging, as no single method currently allows both accurate quantification and spatially resolved visualization. Biochemical assays provide accurate quantification but lack spatial resolution, whereas imaging strategies can perturb membrane organization or cholesterol accessibility.

Here, we describe optimized protocols using fluorescent D4 probes derived from the cholesterol-binding domain of perfringolysin O (D4-mCherry and D4-GFP) to detect, visualize, and quantify cholesterol in biological samples. We detail procedures for probe production, purification, and application, and establish conditions that ensure robust and reproducible labeling of membrane-accessible cholesterol.

By combining fluorescence-based imaging with quantitative analysis, this approach enables the assessment of cholesterol distribution while preserving its native membrane environment. The proposed methodology provides a versatile and reliable framework for studying cholesterol in a wide range of experimental systems.

## Introduction

Lipids are no longer viewed as passive structural elements of the cell membranes but as active molecules that regulate a wild range of signaling processes^1^. Among them, cholesterol occupies a singular position since it accounts for 30-40% of total membrane lipids^2^. Beyond its well-established role in maintaining membrane integrity, cholesterol orchestrates the spatial organization of membranes and controls key signaling pathways. At nanoscale level, cholesterol is a fundamental component of cell membranes, that modulates membrane fluidity and organization. By driving the formation of *lipid rafts*, i.e., dynamic microdomains enriched in cholesterol, sphingolipids, and specific proteins, it plays a key role in signaling processes. Indeed, these rafts serve as functional platforms that compartmentalize cellular processes, facilitating membrane protein sorting, signal transduction, and endocytosis^3,4^. As a results, change in cholesterol content or distribution can disrupt and impair numerous signaling pathways, including those involved in immune responses, neurotransmission, and cell adhesion^5,6^. Consequently, alterations in cholesterol at the membrane can profoundly affect raft-dependent functions and have been implicated in the development of various diseases, from cardiovascular and neurodegenerative disorders to cancers^7,8^.

Despite its fundamental biological importance, studying cholesterol in its native context, at the subcellular level, remains a major challenge. This limitation stems from a fundamental challenge: unlike proteins, cholesterol, is not directly genetically encoded and therefore cannot be tagged using traditional molecular biology tools. As a result, conventional biochemical approaches, including enzymatic colorimetric assays, gas chromatography–mass spectrometry (GC–MS), and high-performance liquid chromatography (HPLC), provide only bulk measurements. Even if these approaches give accurate measurements of cholesterol content, they require cell lysis and / or lipid extraction that abolish spatial information and preclude the analysis subcellular localization of cholesterol.

Conversely, imaging strategies often rely on indirect labeling or sample fixation procedures that can perturb membrane organization and alter cholesterol distribution and accessibility^9^. In addition, preparation steps often involve fixations or detergents that can introduce artifacts, particularly when studying labile lipid–protein interactions. Fluorescent cholesterol analogs, such as BODIPY-cholesterol or dehydroergosterol (DHE), have enabled live-cell imaging but frequently suffer from altered membrane behavior and limited photo-stability, complicating data interpretation^10^. In contrast, protein-based cholesterol probes derived from *perfringolysin O*, such as D4, emerge as a powerful alternative. These probes specifically recognize membrane accessible cholesterol in its native environment without requiring chemical modification of the lipid allowing a live labelling^11,12^.

In this study, we introduce optimized approaches using the D4 probes (D4-mCherry and D4-GFP), derived from the cholesterol-binding domain of *perfringolysin O*^13,14^, to detect, quantify and visualize cholesterol in biological samples. Building on established protocols for the production and purification of these probes^12^, we developed new strategies to enhance their use in the laboratory. Our work demonstrates the potential of the D4 fluorescent probes as robust and versatile tools for investigating cholesterol distribution and quantification on biological samples.

## Material and Methods

### Cell culture

MDA-MB-231 and SH-SY5Y cells from ATCC cells were maintained at 37 °C with 5% CO_2_ in Gibco Dulbecco’s Modified Eagle Medium: Nutrient Mixture F12 (DMEM-F12, GIBCO, Waltham, MA, USA) supplemented with 10% Fetal Bovine Serum (GIBCO, Waltham, MA, USA), 20 mM HEPES (GIBCO, Waltham, MA, USA), 1X Penicillin streptomycin (GIBCO, Waltham, MA, USA).

### D4 plasmids and probes

Plasmids pET28/His6-EGFP-D4 #13961 and pET28/His6-mCherry-D4 #14300 were obtained from the Riken DNA bank (https://dna.brc.riken.jp/en/). Plasmids amplification and probe purification were done following instructions from Wilhelm *et al*^12^.

### Cells treatments

MDA-MD-231 were treated with MβCD (Sigma, St. Louis, MO, USA), 24 h before experimentation at the indicated concentration.

Cells are incubated during 2 h with the D4 probe (mCherry or EGFP) at a final concentration of 3 μg/ml.

### Cell lysis with detergent

Following drug treatment and probe incubation, cells were washed with phosphate-buffered saline (PBS) and lysed in RIPA buffer (150 mM NaCl, 50 mM Tris, pH 7.4, 1% Triton X-100, 0.1% SDS, 1% sodium deoxycholate, all from Sigma, St. Louis, MO, USA) supplemented with Halt phosphatase and protease inhibitor cocktail (Thermo Scientific, Waltham, MA, USA).

### Cell lysis without detergent

Following drug treatment and probe incubation, cells were washed with phosphate-buffered saline (PBS) and lysed in hypertonic buffer (150 mM NaCl, 500 mM sucrose, 50 mM HEPES pH 8.0, 1 M hexylene glycol, all from Sigma, St. Louis, MO, USA) supplemented with Halt phosphatase and protease inhibitor cocktail (Thermo Scientific, Waltham, MA, USA). Cell suspensions were vortexed, freeze overnight at -20 °C and then passed 10 times through a 27G needle. Lysis efficiency was confirmed using trypan blue staining (GIBCO, Waltham, MA, USA) and cell visualization on a hemacytometer (Hausser Scientific, Horsham, PA, USA).

### Protein quantification

Protein concentration was determined using the BCA assay (Thermo Scientific, Waltham, MA, USA) according to the manufacturer’s instructions. Equal amounts of protein were prepared and loaded for Western blot or dot blot analysis.

### Sucrose gradient

HEK-293T cells incubated with D4-EGFP or D4-mCherry for 2h in regular media. Following treatment, cells were washed twice with PBS, collected in PBS, and centrifuged at 2,000 × g for 5 min at room temperature. The resulting pellet was lysed in 500 µL of TKM lysis buffer (0.5 mM Tris, 25 mM KCl, 1 mM EDTA, supplemented with protease and phosphatase inhibitors). One-tenth of the lysate was kept as the total fraction. A total of 2 mg of protein was then loaded onto a discontinuous sucrose gradient (80%, 40% containing the lysate, 36%, and 5%). Samples were centrifuged for 18 h at 171,100 × g. Each 1 ml fraction was collected and precipitated with cold acetone (1:1, acetone:sample) over night at −20 °C, followed by centrifugation at 13,000 × g at 4 °C for 10 min. Each pellet was resuspended in 30 µL of TKM lysis buffer and prepared for Western blot.

### Immunoprecipitation

After probe incubation, cells were washed once with cold PBS, before to be collected in PBS, and centrifuged at 3,500 × g for 5 min at 4 °C. The pellet was washed again with PBS and centrifuged at 3,500 × g for 5 min at 4 °C, before being resuspended in IP lysis buffer (100 mM NaCl (Riedel, #31434), 1 mM MgCl₂ (Merck, #5833), 1% NP-40 (Sigma-Aldrich, #74385), 50 mM Tris-HCl (Fisher Scientific, #ICN819638)) supplemented with Pierce protease inhibitor mini tablets (EDTA-free, Thermo Scientific, Waltham, MA, USA). Lysates were incubated for 1 h at 4 °C under gentle agitation, then centrifuged at 9391 x g for 10 min at 4 °C. Aliquots of 50-175 μg of lysate were immunoprecipitated for 2h at 4 °C with 0.75-1.5 µg of antibody mouse anti-GFP (Novus Biologicals, #9F9.F9). Subsequently, 30-60 μl of Protein G agarose beads (Millipore #P7700 reconstituted in DPBS at 1mg/ml or Pierce™ Protein G Agarose #20398) were added to each lysate and incubated for 18 h at 4 °C. Beads were washed three times with IP buffer 100 mM NaCl, two times with IP buffer 500 mM NaCl, once with IP buffer 100 mM NaCl, once with DPBS and finally resuspended in DPBS before to be prepared for Western Blot. Between washes, beads were centrifugated at 15871 x g for 5 min at 4 °C. Same volume of immunoprecipitated and flow-through fractions (15 ul) was loaded to Western blot gels.

### Electrophoresis and Western Blot

Laemmli sample buffer was added to each sample to obtain a final 1× concentration. For all experiments, 12 μg of total protein were loaded per lane. For co-immunoprecipitation (co-IP) experiment we loaded 15ul of either IP or flowthrough fraction, and 25 µg of total protein for loading control (lysate). Prior to loading onto a NuPAGE™ 4–12% Bis-Tris gel (Invitrogen, Waltham, MA, USA), samples were boiled for 10 min at 70 °C for IP, or, 5 min 90 °C for other purpose, and subsequently transferred to PVDF membrane (co-IP experiment; Amersham ™ Hybond; GE Healthcare, Chicago, IL, USA) activated with 100% methanol (Honeywell™ Riedel de-Haen), or nitrocellulose membrane (GE Healthcare, Chicago, IL, USA). Membranes were blocked in Tris-buffered saline containing 0.1% Tween-20 (TBS-T) and 5% Bovine Serum Albumin (BSA) (Biowest, Nuaillé, France) for at least 30 min at room temperature.

### Dot blot

For all experiments, 15 μg of total proteins were loaded per well. Samples were loaded on the dot blot apparatus (Bio-Rad, Hercules, CA, USA). After Red Ponceau labelling (Thermo Scientific, Waltham, MA, USA), membranes were blocked in Tris-buffered saline containing 0.1% Tween-20 (TBS-T) and 5% BSA for at least 30 min at room temperature.

### Antibodies incubation and Blots revelation

Before antibody incubation, membranes were subjected to Red Ponceau (Thermo Scientific, Waltham, MA, USA) labelling to ensure quality of the transfer. Membranes were then blocked using blocking solution; TBS-T : Tris Buffer Saline, 0.1% Tween supplemented with 2.5% BSA for at least 30 min at room temperature before to be incubated with primary antibodies. The dot blot membranes were incubated in D4-GFP (1.2 ug/ml) overnight at 4 °C where applicable. Primary antibodies used were: chicken α GFP (1:4000 or 1:10 000, Aves Labs #GFP-1010) 1.5 h at room temperature or overnight at 4 °C rabbit α mCherry (1:2000, ProteinTech #26765-1-AP), mouse α Flotillin 1 (1:1000, BD Biosciences #610821) or mouse α Transferrin Receptor (1:1000, Invitrogen #13-6800) overnight at 4 °C. After wash with PBS, membranes were incubated with secondary antibodies incubation (αChicken-HRP, 1:1000, Abcam #ab6877 and αRabbit-HRP, 1:10 000, Cytiva #NA934V) for 1 h at room temperature. Primary and secondary antibodies were diluted in blocking solution. Membranes were washed 3 times with TBS-T after each antibody incubation. Protein detection was performed using the Pierce ECL Western Blotting Substrate (Thermo Scientific, Waltham, MA, USA). Blots were visualized with a G:BOX imaging system (Syngene, Bengaluru, India) using GeneSys software, and band intensity quantification was conducted with ImageJ.

### Blots quantifications

Blots (Western Blots and dot Blots) were quantified with ImageJ (Fiji v.1.54f). Each band was squared, one after another, with the same rectangle’s size (the rectangle was saved using the ROI manager). After that, the rectangle was selected by Analyze > Gels > Select First Lane, then the band was plot by Analyze > Gels > Plot Lanes. Then, the area under the curve was measured with the tool “magic wound”. The value of each band, i.e. each sample, was normalized on the value of the Ponceau’s column. For dot blots, all normalized values were calculated relative to the average of control samples (0 mβCD).

### Immunofluorescence

Before fixation, a homemade anti-cholesterol probe (His₆-mCherry or eGFP-tagged perfringolysin theta toxin D4, respectively from Makino et al., 2017 or Abe et al., 2012) were added to the culture medium at a dilution of 1:250 and incubated for 2 hours at +37°C. After probe incubation, cells were quickly washed with Phosphate Buffer Saline (PBS) and then fixed for 5 minutes with 4% paraformaldehyde (Thermo Scientific, Waltham, MA, USA) in PBS. Cells were washed twice with 1 ml 1x PBS and blocked in staining buffer (SB) containing 2.5% bovine serum albumin, 2.5% normal horse serum, and 5% glycine (all from Sigma, St. Louis, MO, USA) in 1x PBS. Cells were then permeabilized in SB containing 0.3% Triton X-100 (Euromedex, Souffelweyersheim, France) for 30 minutes. Primary antibodies rabbit anti-Caprin-1 polyclonal (Proteintech, # 15112-1-AP), a rat anti-RFP monoclonal antibody for the mCherry (ChromoTek, Planegg, Germany, #5F8) and chicken anti-GFP polyclonal for the eGFP (abcam, Cambridge, UK, #ab13970) were used (1:1000 for all) in IF buffer and incubated overnight at +4°C. The primary antibodies were revealed respectively using a Goat anti-Rabbit AF Plus 405 (For Caprin-1, Invitrogen - #A48254), a Goat anti-Rat Alexa Fluor 568 secondary antibody (For mCherry, Invitrogen - #A32790) and a Goat anti-Chicken AF 488 (For eGFP, Invitrogen - #11039) for 2 hours at room temperature. Finally, after two washes with 1x PBS, cells were mounted using Fluoromount-G™ Mounting Medium (Invitrogen - 00-4958-02).

### Fluorescence image acquisition

Images were scanned on a Zeiss LSM 980 confocal microscope using the ZEN 3.5 software. The images were acquired using a C-Apochromat 40x/1.20 W Korr (stacks of 40-50 images were captured with an interval of 0.2 µm, a voxel size of 207 nm, and an optical zoom of 1.

Laser power and master gain were set based on bright control cells to avoid pixel saturation. All images from a given culture were then acquired using the same acquisition parameters. The raw data were saved at a resolution of 1024x1024 in a 16-bit format. All Z-stacks were maximum intensity projected to facilitate image analysis.

### Image analysis

The analysis was conducted using Fiji v.1.54f. For each probe, a threshold was set to eliminate background noise. Any value below the threshold was marked as "N.A." (not available) to avoid introducing bias in the measurement of mean intensity. For each condition, each value was normalized to the control value of the group (0 mM of MβCD for eGFP or mCherry).

## Results

### Indirect quantification of cholesterol level via Western blot

Our primary goal was to determine whether cholesterol levels could be quantified by Western blot using the D4 probes. To this end, live-cell cholesterol labeling was performed by incubating MDA-MB-231 cells with the D4-GFP or D4-mCherry prior to lysis. After incubation, cells were washed to remove probe excess, lysed, and total protein quantification was performed. As observed, each probe could be detected with a specific antibody against its respective tag, without cross-recognition between probes whereas control cells without any probe incubation showed no signal for either EGFP or mCherry (**Figure 1A**). These results show the specificity of each combination probe/antibody.

**Figure 1:**
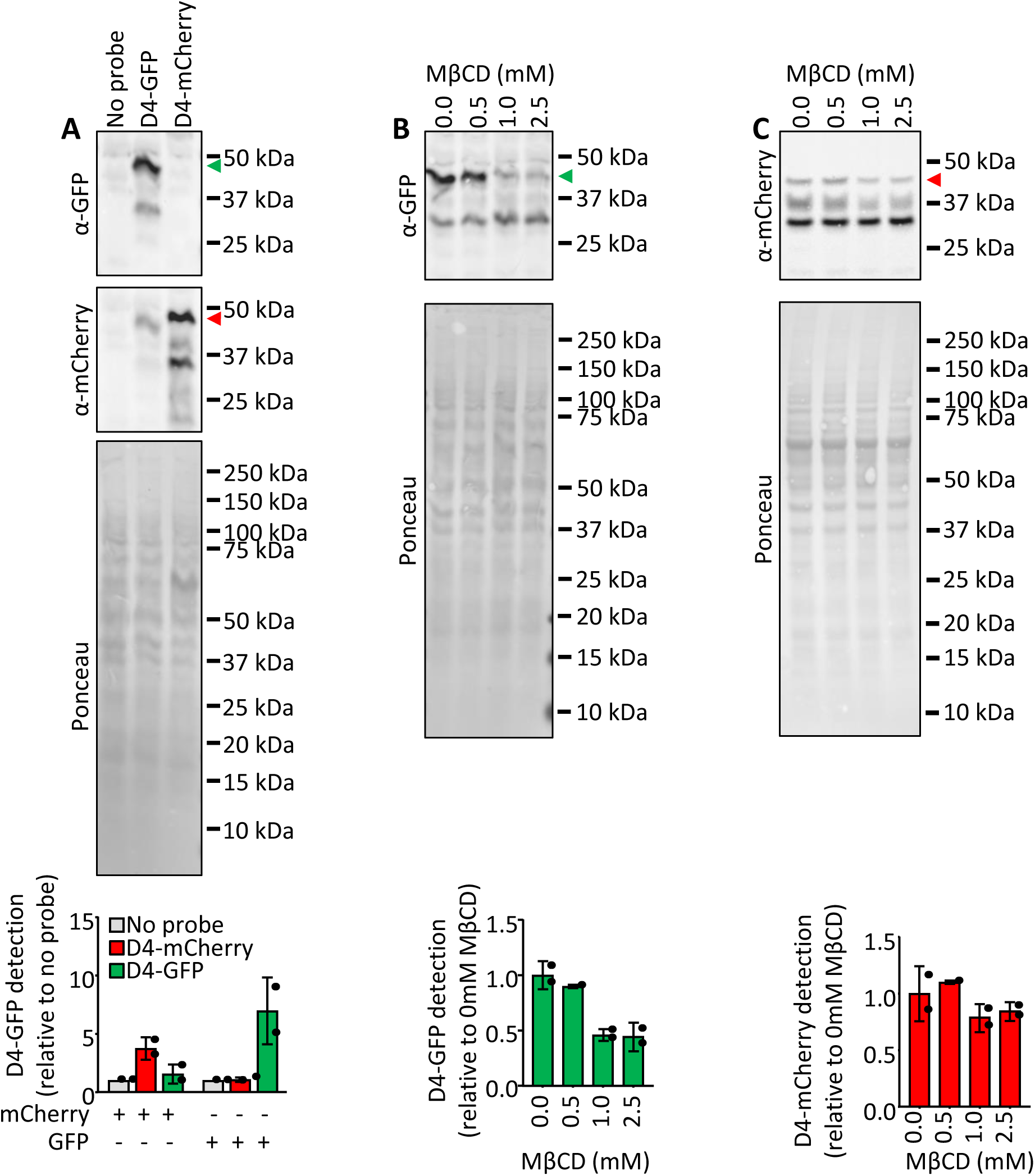
Quantification by Western blot of cholesterol by using D4 probes. **A-C.** MDA-MB-231 cells were incubated with or without D4-GFP or D4-mCherry, lysed before to be subjected to Western blot. Green and red arrows are placed at the respective size of the D4-GFP and D4-mCherry probes. **A.** All three samples, no probe, incubated with D4-GFP or D4-mCherry were revealed with GFP and mCherry antibodies for show specificity. Upper figure; representative blots. Lower figure; quantifications, average + SD. N = 2 independent experiments. **B & C.** Cells were incubated with increased concentration of methyl-beta-cyclodextrin (mβCD) 24 hours before collection. N = 2 for 0.0, 1.0 and 2.5 mM of MβCD and N = 1 for 0.5 mM. Upper figure; representative blots. Lower figure; quantifications, average + SD. N = 2 independent experiments.

To confirm the specificity of cholesterol detection, cells were treated with the well-characterized cholesterol-extracting agent Methyl-β-CycloDextrin (MβCD)^15^. As expected, the probe signal decreased under MβCD treatment but was not completely abolished, as seen on quantification. This likely reflected detection of intracellular cholesterol, that is not affected by MβCD. The concentration-dependent decrease in signal with both probes confirmed the specificity of cholesterol detection by Western blot (**Figure 1B&C**).

### Fast indirect quantification of cholesterol level *via* dot blot

Because Western blot is a powerful but time-consuming technique, we decided to investigate the possibility to perform dot blots to analyze the probes signals. The dot blot assay is faster and simpler, as it does not require electrophoretic separation or transfer steps. This technique also allows high-throughput detection, with up to 96 samples per membrane, and requires less protein amount.

We analyzed in dot blot samples incubated with or without D4-GFP or D4-mCherry. Here we demonstrated that D4-GFP is recognized by the GFP antibody with a clear signal **(Figure 2A)**. While the D4-mCherry presents a GFP intensity similar to the sample without any probe, suggesting antibody background noise (**Figure 2A)**. This highlights the necessity of a negative control in dot blot experiments, and to carefully assess the antibody specificity before starting the dot blot, as the absence of information about protein size can lead to significant background noise. By contrast D4-mCherry is recognized by mCherry antibody in **Figure 2B**, with low background noise for the D4-GFP or the negative control (no probe).

**Figure 2:**
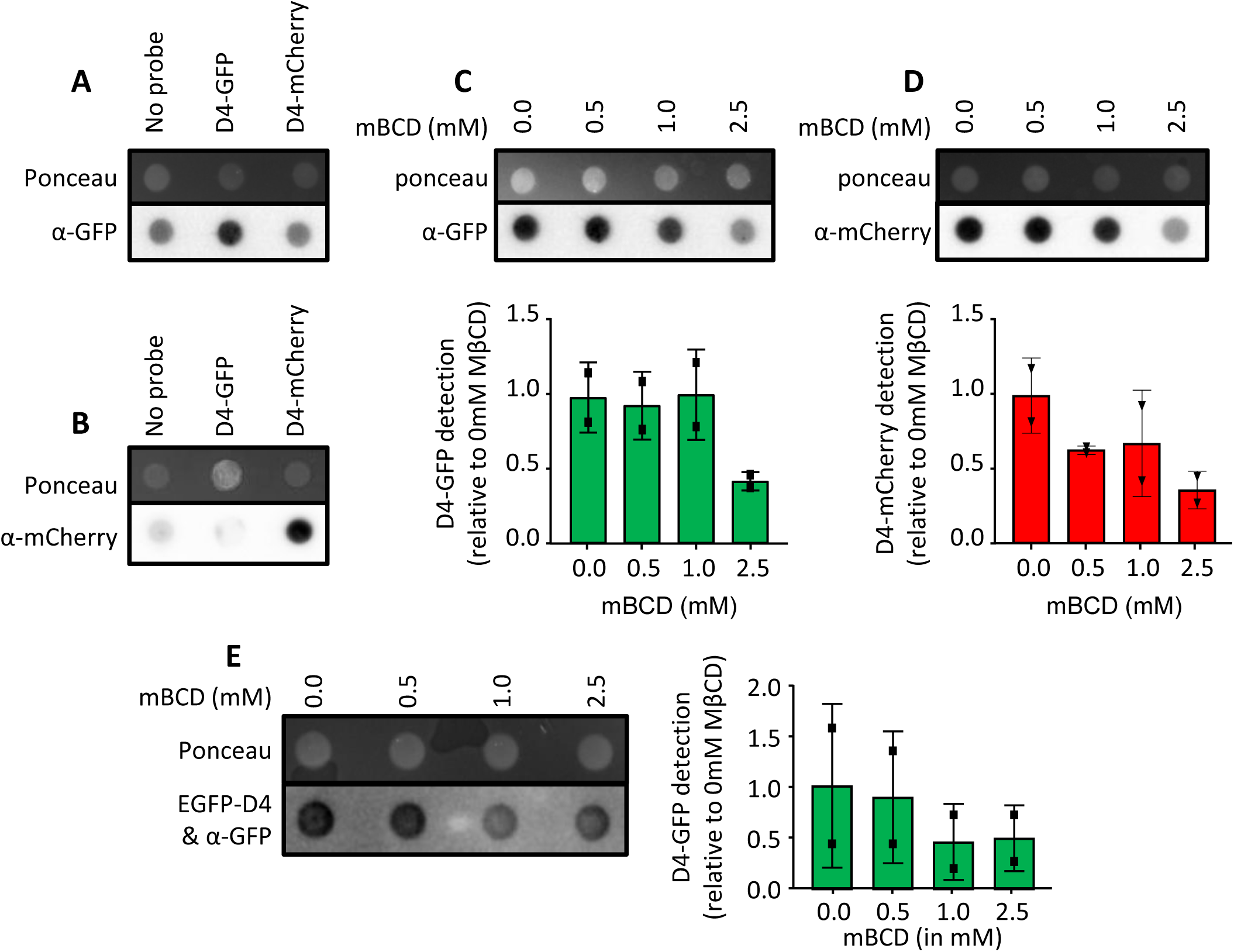
Detection by dot blot of cholesterol by using D4 probes. **A-D.** MDA-MB-231 cells were incubated with or without D4-GFP or D4-mCherry, lysed before to be subjected to dot blot. **A & B.** All three samples, no probe, incubated with D4-GFP or D4-mCherry were revealed with GFP and mCherry antibodies for show specificity. N = 2. **C - E.** Cells were incubated with increased concentration of methyl-beta-cyclodextrin (mβCD) 24 hours before collection. **C & D**. Upper figure; representative blots. Lower figure; quantifications. N = 2. **E.** Cells were lysed without any D4 probe incubation. Cell lysates were subjected to dot blot. Then the blots were incubated first with the D4-GFP probe, then reveal for GFP signal. Left panel; representative blots. Right panel; quantifications. N=2.

Then, the use of MβCD treatment demonstrated the dose-dependent reduction in cholesterol levels for both probes (D4-GFP **Figure 2C**, D4-mCherry **Figure 2D**). This validates the use of D4 probes to monitor cholesterol levels by dot blot analysis. Our next objective was to investigate cholesterol levels by assessing whether the D4 probe could serve as a detection agent post-cell lysis, bypassing the need for live *in vitro* incubation. Cells were lysed with a detergent free hypertonic solution mixed with a freeze-thaw cycle and a mechanical grind. Protein lysates were quantified to load equal amount on dot blot membranes. Then membranes were incubated independently with D4-GFP probe, washed, and revealed thanks to anti-GFP antibody. GFP signal is inversely dose-dependent on the MβCD treatment, consequently, cholesterol levels are assessable via the D4-GFP probe, whether applied in vitro or directly to a blot **(Figure 2E).**

### Visualizing cholesterol via *in cellulo* labelling

Global changes in cholesterol levels can be assessed using Western blot or dot blot techniques; however, these methods do not provide information about the subcellular localization of cholesterol. To address this limitation and visualize cholesterol at the plasma membrane, we investigated the use of fluorescently tagged D4 probes.

Nevertheless, it is important to consider that cell fixation can alter lipid distribution and potentially affect cholesterol labeling. Cholesterol is highly sensitive to organic solvents^16^, which can extract it from membranes during protein fixation. But the use of paraformaldehyde (PFA) also comes with its own limitations. This fixation crosslinks reactive groups on proteins but does not react with cholesterol^17^. As a result, it immobilizes the proteins and therefore can indirectly perturb the surrounding lipid environment, potentially causing cholesterol to redistribute or cluster in non-native membrane regions or making recognition domain not accessible for detection. Consequently, labeling performed after fixation may not accurately reflect the native localization or organization of cholesterol at the plasma membrane. To this regard and to visualize cholesterol at the plasma membrane, we incubated cells with D4 probes prior to fixation, using both available variants: D4-GFP and D4-mCherry.

After probes incubation, cells fixed with cold methanol showed no detectable signal, similar to cells that were not exposed to the probes **(Figure 3A)**. In contrast, cells fixed with PFA post probe incubation displayed a distinct dot-like staining pattern, characteristic of the membrane pattern. Both probes produced comparable staining distributions after paraformaldehyde fixation **(Figure 3B)**. Nevertheless, in cells not treated with D4-GFP, cells displayed a light green auto-fluorescence after PFA fixation but not after methanol fixation **(Figure 3A&B)**. To visualize the cells, we use Caprin-1 a cytoplasmic marker on top of D4-probes labelling for all our immunofluorescence experiments.

**Figure 3:**
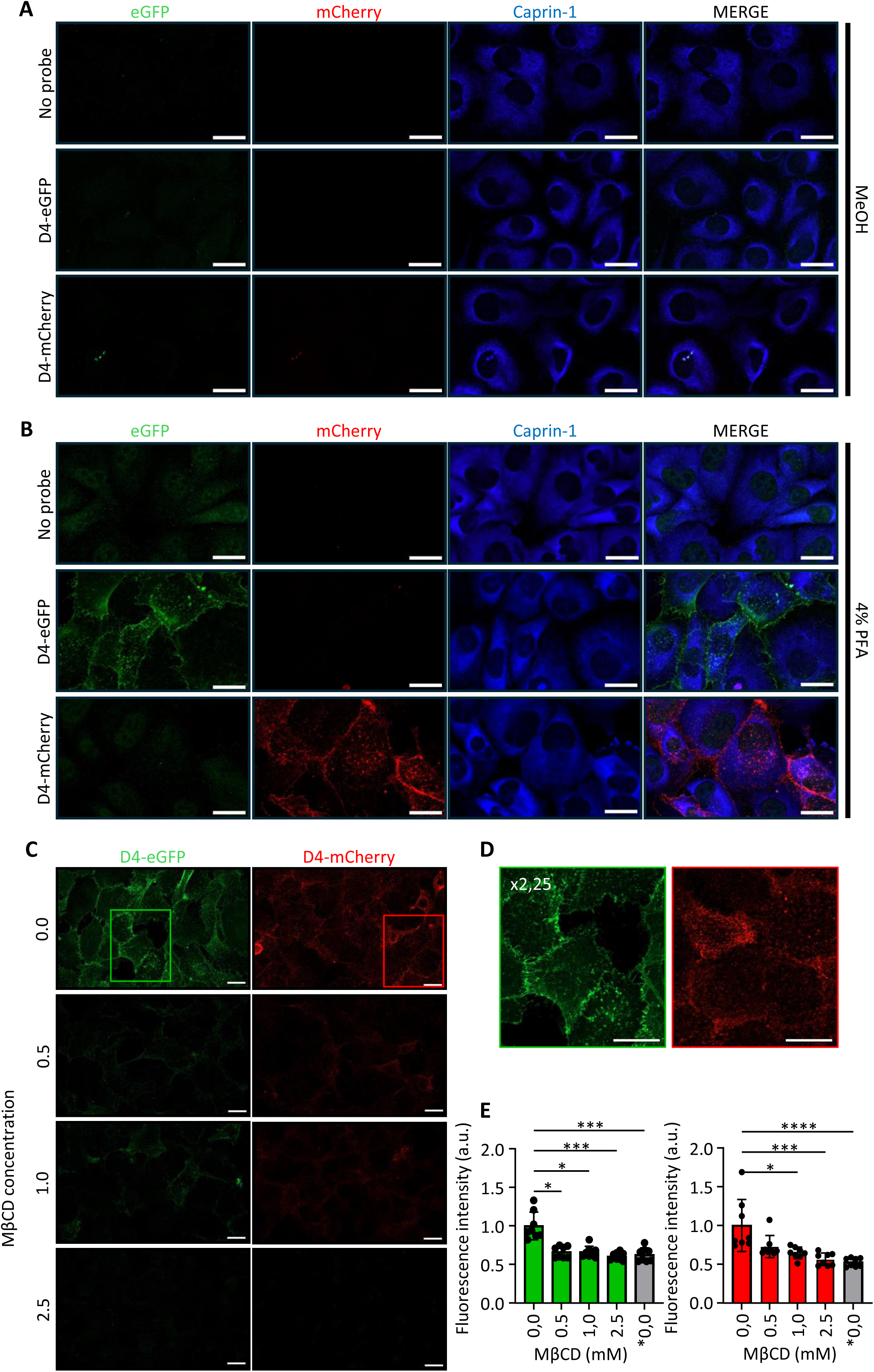
Detection by Immunofluorescence of cholesterol by using D4 probes. **A–D**. Maximum-intensity projections of Z-stack images of immunofluorescence for D4-GFP (green), D4-mCherry (red) and DAPI (blue). Scale bars are all 20 µM **A.** Methanol (MeOH) fixation of cells with or without live incubation with the D4-GFP or D4-mCherry probes **B.** Paraformaldehyde fixation (4% PFA in PBS) of cells with or without live incubation with one of the D4 probes. **C & D.** Cells fixed with 4% PFA and pre-incubated with the different probes or not (0) in the presence of increased concentrations of methyl-β-cyclodextrin (MβCD). **C.** Representative pictures. **D.** Enlargement of the squares represented in C. **E.** Fluorescence intensity quantifications average ± SD. Kruskal–Wallis test was used to compare each MβCD concentration with the corresponding control (CT; 0 mM MβCD) for each probe. Values were normalized to the mean of the control median. Statistical test: * p < 0.05, ** p < 0.01, *** p < 0.001, **** p < 0.0001; ns, not significant. Scale bar: 20 µm.

To further confirm the specificity of the labeling, cells were treated with MβCD. In control cells (untreated with MβCD), both probes produced a clear and distinct membrane-associated pattern, whereas cells exposed to MβCD exhibited little to no labeling, consistent with effective cholesterol depletion. While these observations are qualitative, we developed a quantification protocol based on pixel number and fluorescence intensity to enable comparison between samples **(Figure 3C left)**. As shown in the graph, MβCD efficiently reduces membrane cholesterol even at low concentrations, and the decrease in signal is gradual and concentration dependent **(Figure 3C right)**.

### Detecting the cholesterol in raft via the D4 probes in sucrose gradients

Up to this point, we developed assays to quantify and localize cellular cholesterol. However, given the central role of cholesterol and the growing interest in its membrane-associated functions, we sought to extend our approach. We therefore established an additional assay using D4 probes to investigate cholesterol within lipid rafts. The sucrose gradient flotation assay remains one of the most widely used biochemical techniques to identify lipid raft components. It is useful for studying raft-associated proteins and their distribution.

Lipid rafts can be biochemically isolated using a sucrose gradient assay, which exploits their detergent resistance and low buoyant density. Cells are lysed in cold non-ionic detergents (e.g., Triton X-100 at 4 °C), preserving detergent-resistant membranes enriched in cholesterol and sphingolipids. The lysate is deposed on a discontinuous sucrose gradient and subjected to ultracentrifugation, allowing raft fractions to float to lighter densities.

Collected fractions are subsequently analyzed by immunoblotting for raft markers such as Flotillin 1, and non-raft markers such as Transferrin receptor **(Figure 4)**. Samples with and without probe have very similar profiles for both raft and non-raft markers. Transferrin receptor, a non-raft marker, is distributed relatively homogeneously across fractions 4 to 10. In contrast, the raft marker Flotillin 1 is distributed from fractions 3 to 10, with a stronger signal observed in fractions 3 to 5. The D4-GFP probe co-localizes with Flotillin 1 across fractions 4 to 10, consistent with cholesterol enrichment in lipid rafts. This approach can be used to investigate potential interactions between proteins and cholesterol by assessing their co-localization within the same fractions.

**Figure 4:**
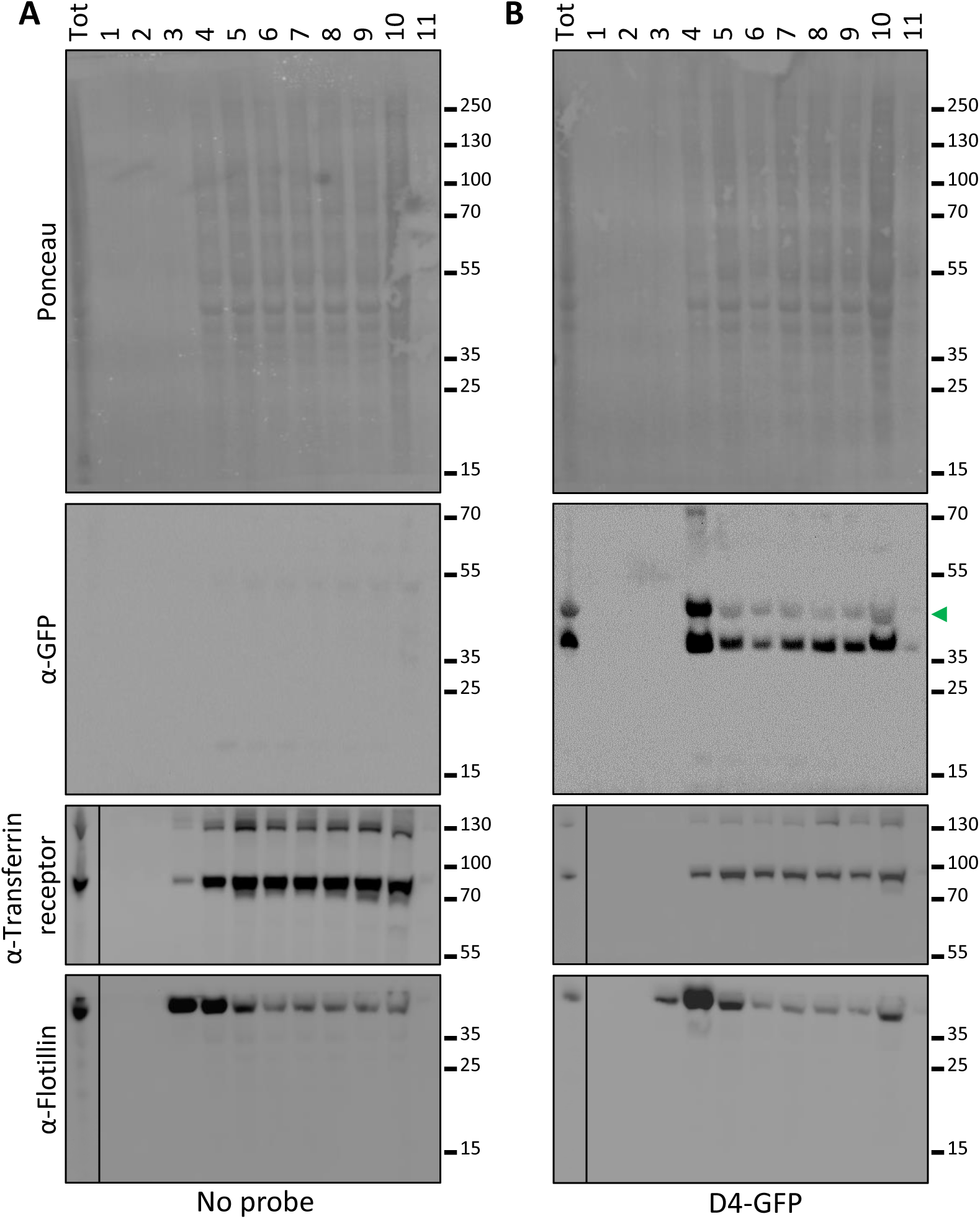
Detection of the D4-GFP in lipid rafts. Sucrose gradient and western blot were performed on MDA-MB-231 cells without (**A.**) or with (**B.)** D4-GFP probe. Flotillin 1 was used as a raft marker and Transferrin Receptor antibody was used as a non-raft marker. An anti-GFP antibody was used to detect the probe. Green arrow is placed at the size of the D4-GFP probe.

### Purifying cholesterol via D4 probes

The last technique we investigated with the D4 probes is the immunoprecipitation. This technique is usually used for the isolation and purification of a specific protein from a complex mixture such as a cell lysate. The method relies on the highly specific interaction between an antibody and its corresponding antigen. Here, we took advantage of the tag, GFP which already have specific antibodies generated for them, to purify the cholesterol bound to the D4-GFP probe.

Cell lysates were incubated with the D4-GFP probe for 2 hours before cell lysis. Then immunoprecipitation was performed using anti-GFP antibody before loading on Western Blot. As observed in **Figure 5**, bands are observed at 45 KDa, the expected size for the D4-GFP probe. In the IP lane, we observed a band at the expected size that is not present in the control IP. In contrary to the flow through, were we observe a band in the negative control and not in the GFP IP relative control. This shows that we successfully immunoprecipitated the probes. This provides the proof of concept that the probes could immuno-precipitated and potentially be used to follow interactions between cholesterol and proteins.

**Figure 5:**
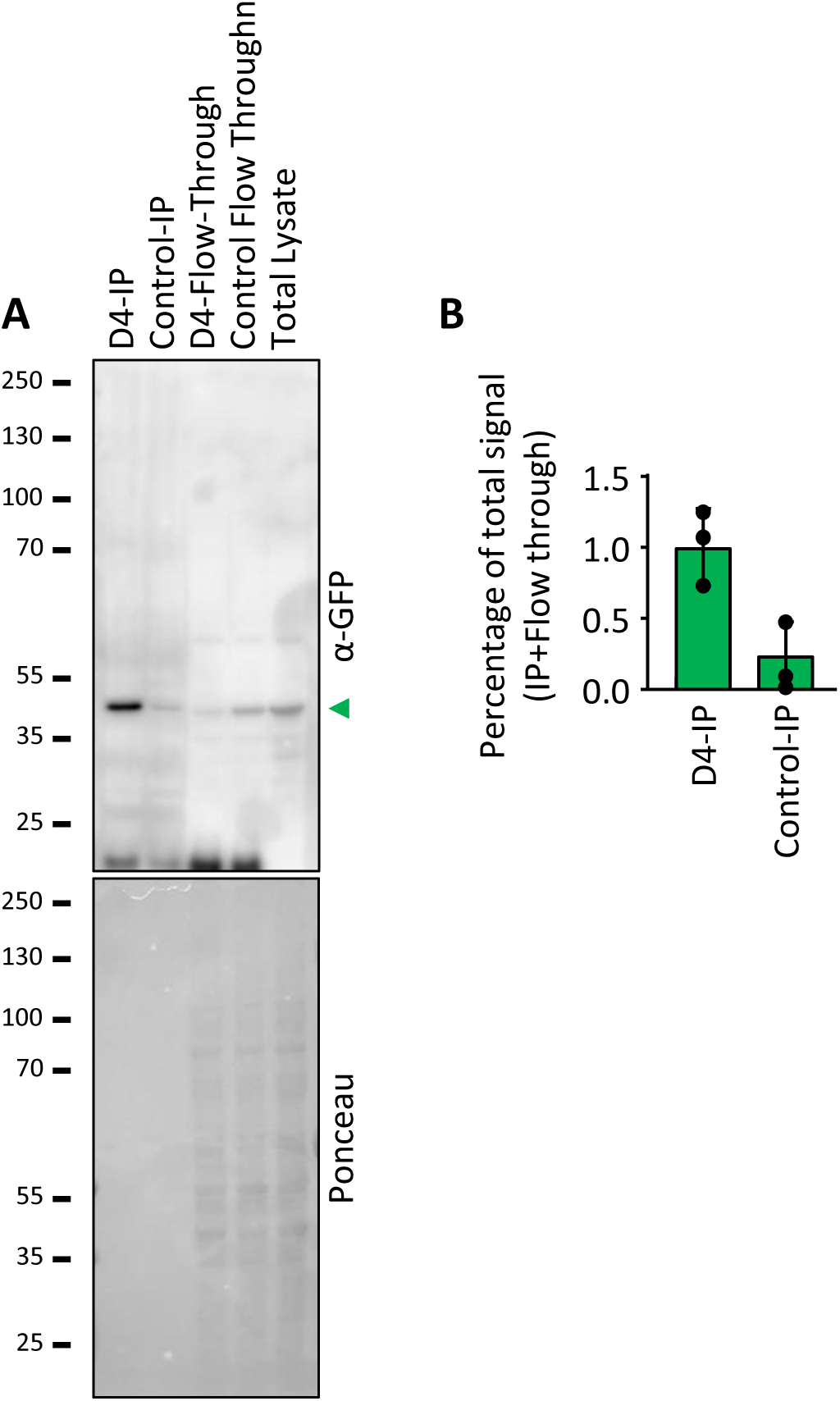
Immunoprecipitation of the D4-GFP probe. **A & B.** MDA-MB-231 cells were incubated with or without D4-GFP, lysed and subjected to immunoprecipitation using and anti-GFP antibody (or Control IP). IP and first flow-through fractions were loaded on Western blot along with the total lysate (N=3). **A.** Representation blots **B.** Quantifications

## Discussion

In recent years, the field of lipid biology has received increasing attention, as lipids are now recognized not only as structural components of cell membranes but also as key regulators of signaling, trafficking, and metabolic processes. Among them, cholesterol occupies a unique and central position. Despite its biological importance, studying cholesterol in native conditions remains challenging due to its small, mostly hydrophobic structure and minimal polar headgroup, which prevent the generation of antibodies. Consequently, researchers have had to develop alternative strategies to detect, quantify, and visualize cholesterol in biological systems.

In this study, we optimized and expanded the use of D4 fluorescent probes, derived from the cholesterol-binding domain of *perfringolysin O*^11,12^, to provide versatile tools for cholesterol detection and quantification. By combining quantitative (Western blot and dot blot) and qualitative (immunofluorescence) assays, as well as immunoprecipitation-based purification, we established complementary approaches that together offer a comprehensive overview of cholesterol levels and localization within cells.

The Western blot and dot blot assays demonstrate that D4 probes can be used for specific detection of cholesterol-associated signals in cell lysates, with the dot blot approach offering a rapid, high-throughput alternative that conserves sample and time. The consistent decrease in D4 signal upon cholesterol depletion with methyl-β-cyclodextrin confirms the specificity of the detection and validates the D4 probe as a reliable indirect reporter of cholesterol content.

The immunofluorescence experiments further highlight the potential of D4 probes to visualize cholesterol at the plasma membrane in living cells. Importantly, our results emphasize that fixation conditions strongly influence the accuracy of cholesterol labeling: methanol fixation completely abolishes the signal, while paraformaldehyde preserves the membrane-associated pattern. These findings align with previous observations that fixation and solvent exposure can alter lipid distribution or detection, underlining the necessity of optimizing labeling conditions when working with lipid-binding probes. In future applications, this immunofluorescence protocol could enable co-localization studies of cholesterol with specific proteins, thereby providing insights into membrane organization and raft-associated processes^18^.

Finally, our immunoprecipitation experiments provide a proof of concept that D4 probes can be used not only for visualization but also for the isolation of cholesterol-associated complexes. The successful co-purification of cholesterol with D4 probes supports the idea that these tools could be applied to identify cholesterol–protein interactions under native conditions. This opens new possibilities for exploring the molecular partners of cholesterol and understanding its functional roles in diverse cellular contexts.

## Conclusions

Overall, this study establishes and refines practical protocols for the use of D4-based cholesterol probes across different experimental applications. The integration of these quantitative and qualitative approaches provides robust cross-validation of probe specificity and extends the toolbox available for cholesterol research. Future studies could combine these methods with advanced imaging techniques, such as super-resolution microscopy or live-cell imaging, to further unravel the spatiotemporal dynamics of cholesterol and its involvement in cellular signaling and membrane organization.

## List of abbreviations

BODIPY: Boron–dipyrromethene
Co-IP: Co-immunoprecipitation
D4: Domain 4
DHE: Dehydroergosterol
DMEM/F-12: Dulbecco’s modified Eagle medium and Ham’s F-12 medium
DPBS: Dulbecco’s Phosphate-Buffered Saline
EDTA: Ethylenediaminetetraacetic acid
EGFP: Enhanced Green Fluorescent Protein
GC-MS: Gas chromatography–mass spectrometry
GPI: Glycosylphosphatidylinositol
HEPES: 2-(4-(2-hydroxyethyl)-1-piperazinyl)-ethanesulfonic acid
HPLC: High-performance liquid chromatography
MβCD: Methyl-β-CycloDextrin
MDA-MB-231: M D Anderson - Metastatic Breast – 231
ROI: Region of interest
PBS: Phosphate Buffer Saline
SB: Staining buffer
TBS: Tris Buffer Saline
TBS-T: Tris Buffer Saline Tween

## Declarations

### Ethics approval and consent to participate

Not applicable

### Consent for publication

Not applicable

### Availability of data and materials

All data generated or analyzed during this study are included in this published article

### Competing interests

There is no competitive interest

### Funding

This work is supported by the Research Council of Finland (grants 333096, 335956 and 358122) and by the Sigrid Jusélius Foundation to CDS. The HiLIFE fellow program and the Neuroscience Center (University of Helsinki) supports CDS as a young group leader.

CK has received support from Finnish Epilepsy Research Foundation (Epilepsiatutkimussäätiö) and Finnish Instrumentarium foundation (Instrumentariumin Tiedesäätiön).

APLC has received support from Epilepsiatutkimussäätiö and The Finnish National Agency for Education (EDUFI Fellowship OPH-682-2024).

EB has received support from Finnish Epilepsy Research Foundation (Epilepsiatutkimussäätiö).

### Authors’ contributions

CDS had the original idea, provided funding and administrated the project. AA planned and supervised the experimental part. AA & AT produced and purified the D4 probes. AA, EB, CK, APLC and AT performed and adjusted the experiments. EB, CK and APC analyzed the results. AA, EB, CK and APLC produced the figures. AA and CDS wrote the manuscript. All the authors corrected and approved the manuscript.

## Acknowledgements

We would like to thank the HiLIFE Neuroscience Center at the University of Helsinki for providing access to the cell culture and microscopy facilities, as well as to the equipment, including the Leica LSM 980 microscope and the Syngene G:BOX imaging system. We are also grateful to Dr. Eero Castrén for granting access to the dot blot apparatus, and to Seija Lågas for her excellent technical assistance and consistently cheerful support. We also thank the RIKEN BRC who provided the pET28/His6-EGFP-D4 (cat#RDB13961) and pET28/His6-mCherry-D4 (cat#RDB14300) constructs.

## Authors’ information

EB, CK and APLC PhD students

AT: Summer student

AA: Research Coordinator

CDS: Principal Investigator

## Footnotes

Not applicable

## Notes

### Competing Interest Statement

The authors have declared no competing interest.

